# Under the radar: differential responses of bed bugs to an entomopathogen, environmental bacteria, and a human pathogen

**DOI:** 10.1101/2024.02.28.582586

**Authors:** Hunter K. Walt, Aline Bronzato-Badial, Sophie E. Maedo, Joseph A. Hinton, Jonas G. King, Jose E. Pietri, Federico G. Hoffmann

## Abstract

**Background:** Bed bugs (Hemiptera: Cimicidae) are a widely distributed, obligately blood-feeding insect, but they have never been linked to pathogen transmission in humans. Most other hematophagous insects that frequently bite humans transmit pathogens, and it is unclear why bed bugs do not. One hypothesis is that bed bugs have evolved a highly robust immune system because their mating system, traumatic insemination, exposes females to consistent wounding and bacterial infections. Although this has been proposed, very little is known about the bed bug immune system and how bed bugs respond to microbial challenges. Understanding the bed bug immune system could give insight to why bed bugs are not known to transmit disease and under what circumstances they could, while also facilitating biological control efforts involving microbes.

**Methods:** To investigate the immune response of bed bugs to bacterial challenges, we exposed female bed bugs to three bacterial challenges. 1.) *Pseudomonas fluorescens*, an entomopathogen known to have harmful effects to bed bugs, 2.) bacteria cultured from a bed bug enclosure likely encountered during traumatic insemination, and 3.) *Borrelia duttoni*, a human vector-borne pathogen that causes relapsing fever. We compared the transcriptomes of infected bed bugs with uninfected bed bugs, focusing on immune-related genes. We also conducted phylogenetic analyses to understand patterns of gene duplication and function of potentially immune-related genes.

**Results:** We found many known immune effector genes upregulated in response to *P. fluorescens* and traumatic insemination-associated bacteria, but interestingly, not in response to *B. duttoni*. Furthermore, we found significant overlap in the genes differentially expressed in response to *P. fluorescens* and the traumatic insemination associated bacteria, and between *P. fluorescens* and *B. duttoni*, but no significant overlap between traumatic insemination bacteria and *B. duttoni*. We also show that bed bug diptericin-like antimicrobial peptides underwent a lineage-specific gene duplication, and that they may have further functional specialization. Finally, we identify previously overlooked candidates for future study of immune function in bed bugs, including some putative cuticle-associated genes, a laccase-like gene, and a mucin-like gene.

**Conclusions:** By taking comprehensive transcriptomic approach, our study is an important step in understanding how bed bugs respond to diverse immune challenges.

## Background

Bed bugs (Hemiptera: Cimicidae) are important hematophagous pests and public health concerns, as their bites can cause allergic reactions and infestations are psychologically damaging and notoriously hard to control, but they are not known to transmit pathogens outside of a laboratory setting [1–4]. This is interesting as most blood-sucking insects that routinely feed on humans are associated with pathogen transmission. There have been many hypotheses addressing why bed bugs do not transmit pathogens, but few have been tested [4–6]. One of these hypotheses suggests that bed bugs do not transmit disease because they have evolved a particularly robust immune system due to consistent wounding and bacterial exposure associated with their mating system, traumatic insemination [3–5,7]. During traumatic insemination, a male bed bug bypasses the female’s genital orifice and directly pierces her abdomen with a sharp intromittent organ to release sperm directly into the hemolymph [8]. Interestingly, female bed bugs have evolved a novel organ called the spermalege, likely in response to sexual conflict [9–12]. The spermalege is known to reduce the physical costs of traumatic insemination because the cuticular portion (ectospermalege) is rich in resilin, an elastic protein that allows the cuticle to extend and contract resiliently [13,14]. Along with this, the spermalege is filled with cells called hemocytes, some of which control cellular immune responses in insects via encapsulation, nodulation, and phagocytosis of pathogens [8,15–17]. This suggests that the spermalege may mitigate bacterial infection associated with wounding from traumatic insemination [11,18]. Consistent with this, another study found that antimicrobial peptide (AMP) transcripts were highly expressed in the bed bug spermalege [19].

Although some studies have measured fitness effects of bed bugs exposed to various bacteria [20], the molecular immune response of bed bugs to bacteria has not been well characterized. The insect molecular (or humoral) immune response is activated when host-derived pattern recognition receptors (PRRs) bind to pathogen-associated molecular patterns (PAMPs) such as lipopolysaccharide or peptidoglycan from bacteria [21]. When PAMPs bind to PRRs, signaling pathways are induced that produce antimicrobial effectors such as AMPs or phenoloxidases [21,22]. Annotations of the bed bug genome showed that, similar to other hemipteran insects, their immune repertoire is lacking in some canonical insect immune-related genes such as parts of the Imd pathway [14,23]. Along with this, bed bugs have a relatively low number of identifiable AMPs, with genes encoding three defensin AMPs and two diptericin-like putative AMPs [14,24]. Previous studies found that the three defensin AMPs are upregulated in response to various bacterial challenges, and another study found the two diptericin-like genes expressed in bed bug hemocytes [17,24]. Along with this, several studies have shown that bed bug extracts have antimicrobial properties [25–27]. Furthermore, many studies have assessed the competence of bed bugs as vectors of various human pathogens, but most of them did not measure the immunological response of bed bugs to these pathogens [28–31]. One study found evidence that the human pathogen *Borrelia recurrentis* is detected by hemocytes in culture, but did not measure humoral responses [32].

Only one study has considered the whole transcriptomic response of bed bugs to immune insult, but this study included only a single stimulus (*E. coli*) and focused only on hemocytes in an *in vitro* context [17]. To fill this gap, we investigated the transcriptomic response of intact bed bugs to three bacterial challenges. First, we exposed bed bugs to a known entomopathogen and potential biological control agent, *Pseudomonas fluorescens,* via intrathoracic prick. *P. fluorescens* has known adverse fitness effects to bed bugs [20], so we predict it will elicit the differential expression of immune-related genes and serve as a baseline understanding for the transcriptomic response to immune challenge. Second, we exposed bed bugs to a bacterial culture grown from their enclosure environment, which allows us to assess the transcriptomic response to bacteria that female bed bugs are likely exposed to during traumatic insemination. Finally, we challenged bed bugs with *Borellia duttoni* via a contaminated blood meal. *B. duttoni* is pathogenic to humans as the causative agent of tick-borne relapsing fever. Bed bugs have been shown to be competent vectors of the closely related spirochete *B. recurrentis* [29] in a laboratory setting, and a previous study claimed *B. duttoni* survived for at least 150 days in bed bugs, making it an interesting model to examine the immune response of bed bugs to vector-borne pathogens [33]. Our treatments were designed to mimic the way that bed bugs may encounter the treatment bacteria naturally, with the entomopathogen and the environmental bacteria being inoculated through a cuticle prick and the human pathogen being administered via a blood meal. All the treatments were paired with corresponding controls, and we sequenced the transcriptomes of individual adult females. We compared the transcriptomes of each treatment to their controls, taking special interest in immune-related transcripts to shed light on the molecular response to immune insult in bed bugs.

## Methods

### Pseudomonas fluorescens infections

Infection experiments with entomopathogenic bacteria were conducted in the Cincinnati field strain of *Cimex lectularius* maintained in the Pietri Lab insectary at the University of South Dakota. Bed bug colonies were kept in plastic jars containing corrugated cardboard harborages at 28 +/- 1° C and 60-70% relative humidity with a photoperiod of 12:12 hours (L:D) and fed once per week using mechanically defibrinated rabbit blood (Hemostat Laboratories, Dixon, CA) once per week using a Hemotek membrane feeding system (Hemotek LTD, Blackburn, UK). Female bed bugs that had not been fed for one week were individually challenged with *Pseudomonas fluorescens* via thoracic prick as previously described [20]. In brief, bacterial cultures were grown overnight in LB media at room temperature and subsequently diluted to a concentration of 10^8^ CFU/ml. Insects were anesthetized on ice, then, to carry out infections, a sterile pin was dipped into the bacterial solution and directly inoculated into the ventral thorax by gently piercing the cuticle. Following inoculation, the insects were kept at 37°C for 6 hours. At 6 hours post-infection, the bed bugs were individually placed into tubes containing TRIzol reagent (ThermoFisher, Waltham, MA) and frozen at −80°C for extraction of RNA. Bed bugs wounded with sterilized pins served as controls for these experiments.

### Borrelia duttoni infections

Bed bugs were maintained in the same way as with the *P. fluorescens* infections. Female bed bugs were starved for one week prior to infection with *Borrelia duttoni*. Stocks of *B. duttoni* (NIH Rocky Mountain Laboratories, Hamilton, MT) were grown in BSK-H medium with 6% rabbit serum (Sigma Aldrich, St. Louis, MO) until the culture reached log phase (∼4 days). To prepare an infectious blood meal, the concentration of spirochetes in the culture was measured by counting on a hemocytometer and adjusted to 10^5^/ml using aseptic, defibrinated rabbit blood (Hemostat Laboratories). This blood meal was provisioned to bed bugs using a Hemotek feeder until they fully engorged. Following feeding, the insects were kept at 28° C for 24 hours. At 24 hours post infection, a total of three female insects were individually placed into tubes of TRIzol reagent (ThermoFisher, Waltham, MA) and frozen at −80° C for extraction of RNA. Bed bugs fed aseptic, defibrinated rabbit blood served as controls for these experiments.

### Traumatic insemination bacteria infections

Traumatic Insemination bacteria (TI bacteria) experiments were conducted in the Harlan strain of *C. lectularius* maintained in the King Lab at Mississippi State University, Starkville, MS. Bed bug colonies were kept in glass jars containing corrugated cardboard paper at 28 +/- 1° C, 60-70% humidity, and photoperiod of 12:12 hours (L:D). Colonies were fed once a week on a human volunteer. Prior to TI bacterial infection, female bed bugs were identified during the nymph stage and maintained separated until the adult stage. Once in the adult stage, bed bugs were starved for one week before TI bacterial infections and were individually challenged with a bacterial solution via intrathoracic prick. The bacterial solution was obtained from a paper swab from the glass jars where bed bugs colonies were maintained. The swab was kept overnight in liquid Luria Broth media at 37°C and rotation at 250 rpm. Insects were anesthetized on ice, and TI bacterial infection was carried out using a sterile pin dipped into the bacterial solution and directly inoculated into the ventral thorax by gently piercing the cuticle. Following inoculation, female insects were kept separated in different vials at 28°C for 24 hours. At 24 hours post-infection, insects were individually placed into tubes of RNAzol RT reagent (Molecular Research Center, Cincinnati, OH) and frozen at −80°C for total RNA isolation. Starved and non-wounded (naive) bed bugs were used as controls for these experiments.

### RNA Extraction

For insects used in the TI bacterial infection experiments, total RNA was extracted from individual bed bugs using RNAzol RT reagent (Molecular Research Center, Cincinnati, OH). For bed bugs challenged with *P. fluorescens* and *B. duttoni*, total RNA was extracted from individual insects using Trizol reagent (Sigma-Aldrich, Burlington, MA).

### RNA-seq

To understand global changes in the bed bug transcriptome under the various challenges, we sequenced the RNA of 18 individual female bed bugs. For each treatment (*P. fluorescens*, *B. duttoni*, and TI bacteria), we took isolated RNA from three treated females, and three control females. Illumina library prep was conducted by Novogene with poly-A enrichment for mRNA, and cDNA was sequenced on a NovaSeq 6000 instrument generating 150 base pair paired-end reads. We deposited raw sequencing reads into NCBI’s sequence read archive under the BioProject accession PRJNA1079782.

### Differential Gene Expression Analysis

Reads were quality inspected by FastQC v0.11.5 [34] and quality trimmed using trimmomatic v. 0.39 [35]. Transcript abundance was measured by quasi-mapping using Salmon v. 1.4.0 [36] with the reads being pseudo-aligned to the *Cimex lectularius* reference genome (GCF_000648675.2) and transcriptome. Data was imported to R using tximport [37]. Differential gene expression analyses were conducted using BioConductor’s DEseq2 package [38], separately analyzing each paired dataset. To test for functional enrichment in the differentially expressed genes, we annotated the *C. lectularius* proteome with Gene ontology (GO) terms using eggNOG mapper v2.1 .9 in Diamond mode [39–41]. To test for enrichment of GO terms in the differentially expressed bed bug genes, we used topGO v2.50.0 [42] in R, using Fisher’s classic test, only considering differentially expressed genes with an adjusted p-value ≤ 0.05. To determine if the number shared, differentially expressed genes were more than you would expect at random, we used Fisher’s Exact test for count data in R [43]. For each treatment pair, we used the number of genes that were expressed in both treatments as the sample.

### Phylogenetic Analysis

To investigate phylogenetic relationships between genes that were differentially expressed in response to immune challenges, we sampled putative homologous genes using BLASTp, against NCBI’s nr protein database, and tBLASTn against NCBI’s Transcriptome Shotgun Assembly database (TSA) and NCBI’s Whole genome Shotgun database (WGS). We obtained coding sequences of the WGS hits using efetch. For the TSA hits, coding sequences of the putative homologs were obtained using the getorf tool from EMBOSS (https://www.ebi.ac.uk/Tools/emboss/). All coding sequences were conceptually translated to amino acid sequences using the transeq tool from EMBOSS. The protein sequences were aligned with MAFFT v7.490 using the L-ins-I algorithm, and the resulting alignment was backtranslated using PAL2NAL [44]. Maximum likelihood phylogenetic trees were built using IQ-TREE v2.0.7 [45], incorporating the ModelFinder algorithm to detect the best substitution model [46]. Branch support was estimated using ultrafast bootstrap with 1000 replicates, the Shimodaira-Hasegawa-like approximate likelihood ratio test (SH-aLRT) with 1000 replicates, and the aBayes test, but only ultrafast bootstrap values were considered for our phylogenetic analyses [47–50].

For the diptericin-like gene phylogeny, we identified putative homologs using BLASTP against NCBI’s non-redundant protein database, keeping only the longest isoform for the phylogeny. We added Diptericin A and Diptericin B from *Drosophila melanogaster*, even though they were too diverged to be detected by BLAST. We followed the same procedure described earlier for phylogenetic analysis, excluding backtranslation step, as we used the protein alignment to build the phylogenetic tree. The clade of diptericin-like genes from Odonata was used as the outgroup, as they belong to the earliest branching clade based off a previous phylogenomic study [51].

## Results

We exposed female bed bugs with three different bacterial challenges: 1.) the entomopathogen *Pseudomonas fluorescens* 2.) a bacterial culture grown from a bed bug enclosure which we refer to as TI bacteria, as they are likely the bacteria that bedbugs would be exposed to during traumatic insemination, and 3.) *Borrelia duttoni*, a human vector-borne pathogen. After differential expression analysis, we found upregulation of five immune effector genes in response to *P. fluorescens* and the TI bacteria, but interestingly, not in response to *B. duttoni* (**Fig. 1, Additional File 1: Fig. S1**). These five effector genes included two defensin-like genes, two diptericin-like genes, and one lysozyme-like gene. For clarity, we hereon refer to the defensin-like genes LOC106661791 and LOC106661792 as Cl-defensin A and Cl-defensin B, respectively. Along with this, we will refer to the lysozyme-like gene, LOC106663700, as Cl-lysozyme, and the two diptericin-like genes, LOC112127760 and LOC106664366, as Cl-prolixicin A and Cl-prolixicin-B, respectively. We refer to the diptericin-like genes as “Cl-prolixins” because our phylogenetic analysis revealed that they are more closely related to the prolixicin AMP of *Rhodnius prolixus* than to diptericin. Finally, we examined shared responses between all treatments which identified potentially important genes involved in bed bug immunity that have been neglected in past studies.

**Fig. 1.**
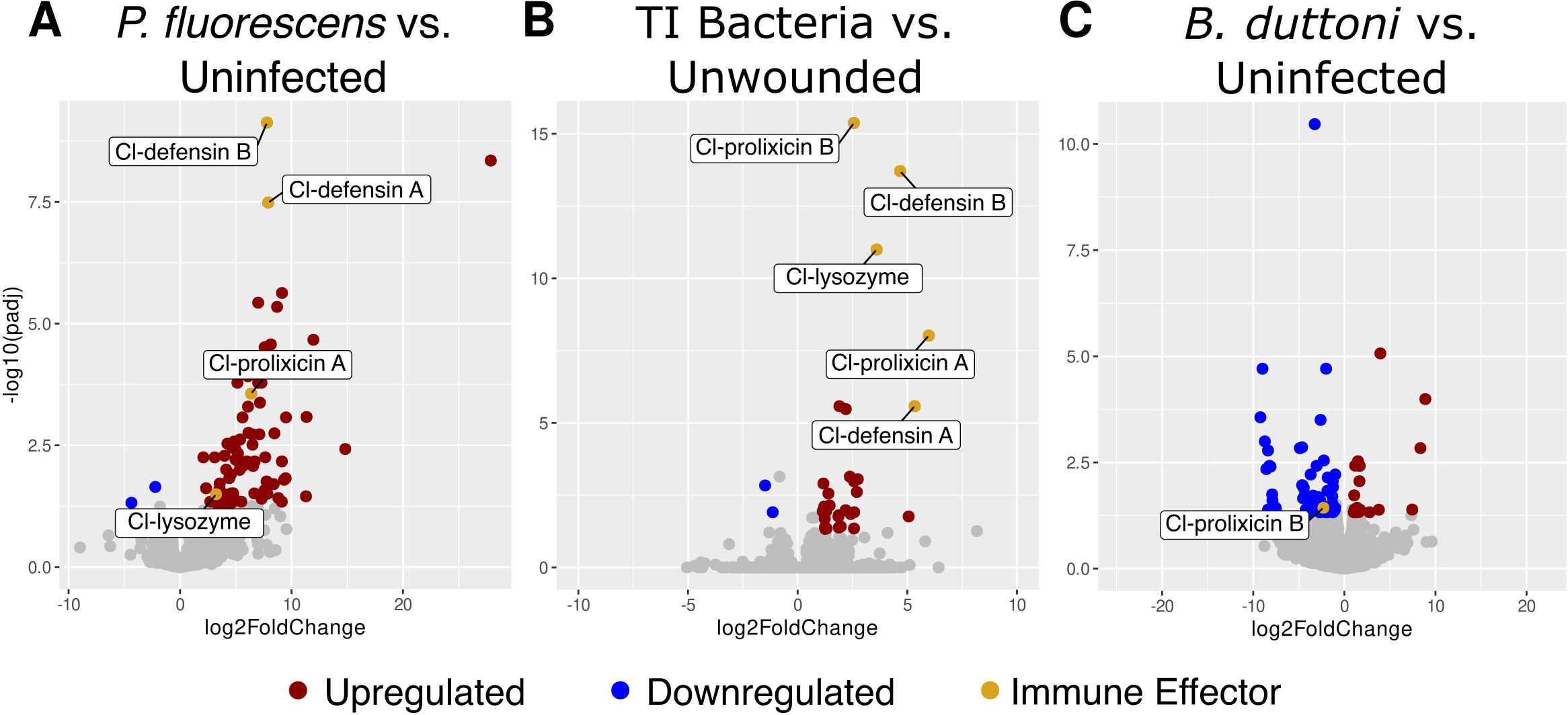
Global changes in bed bug gene expression under varying immune challenges. **(A)** Changes in bed bug gene expression when infected with the entomopathogen *P. fluorescens* via thoracic prick vs. sterile thoracic prick. (**B)** Changes in bed bug gene expression when infected with the human pathogen *B. duttoni* via an infected blood meal vs. a sterile blood meal**. (C**) Changes in bed bug gene expression when traumatic insemination is simulated using a contaminated needle prick vs. bed bugs that were unwounded.

### Infection with Pseudomonas fluorescens

When infected with *P. fluorescens*, a known entomopathogen, we found 79 genes upregulated, and two genes downregulated in adult, female bed bugs (**Fig. 1A**). As expected, antimicrobial effector genes were upregulated in response to this treatment, indicating that *P. fluorescens* is detected by the bed bug humoral immune system. The antimicrobial effectors included Cl-defensin A and Cl-defensin B, Cl-lysozyme, and Cl-prolixicin A. Along with microbial defense responses, GO terms for genes associated with the insect cuticle were highly enriched in this treatment (**Table 1**).

**Table 1.** Top 10 significantly enriched biological process GO terms in the genes differentially expressed in response to infection with the entomopathogenic bacteria *P. fluorescens*.

| GO ID | Term | P value |
| --- | --- | --- |
| GO:0040003 | Chitin-based cuticle development | 1.3e-25 |
| GO:0042335 | Cuticle development | 3.6e-23 |
| GO:1901073 | Glucosamine-containing compound biosynthetic process | 9.6e-09 |
| GO:0046349 | Amino sugar biosynthetic process | 8.9e-08 |
| GO:1901971 | Glucosamine-containing compound metabolic process | 1.9e-07 |
| GO:0006040 | Amino sugar metabolic process | 6.1e-07 |
| GO:0006030 | Chitin metabolic process | 1.0e-06 |
| GO:0006031 | Chitin biosynthetic process | 1.4e-06 |
| GO:0050830 | Defense response to Gram-positive bacterium | 0.00017 |
| GO:0006022 | Aminoglycan metabolic process | 0.00036 |

The *P. fluorescens* treatment had the two most highly differentially expressed genes in our study. These genes were LOC106661042 (acanthoscurrin-like), which had a log2 fold change of 27.88 and LOC1066641041 (glycine-rich protein DOT1-like), which had a log2 fold change of 14.82. They encode for highly similar proteins (amino acid percent identity: 94.27%) and occur in tandem in the *C. lectularius* genome. To assess their evolutionary relationship, we built a phylogenetic tree with these genes along with a selection of putative homologs identified from BLAST and confirmed that they are indeed paralogous (**Fig. 2**). The two paralogs group in a moderately supported clade (bootstrap = 87) with other heteropteran insects including two species of kissing bugs, two species of stink bugs, and *Orius laevigatus* (predatory minute pirate bug) (**Fig. 2**). Of note, the two *Triatoma infestans* sequences included in our phylogeny are likely the same gene but are derived from two independent transcriptome assemblies. Database searches and our phylogeny indicate that this gene family is only present in the Heteroptera and underwent a duplication in the bed bug lineage (**Fig. 2**). Despite these genes being paralogs; their functional annotations are different on NCBI. This, along with no identifiable conserved protein domains, leaves the role of their gene products unknown, although their high expression in response to an immune challenge suggests they may play a role in responding to immune insult.

**Fig. 2.**
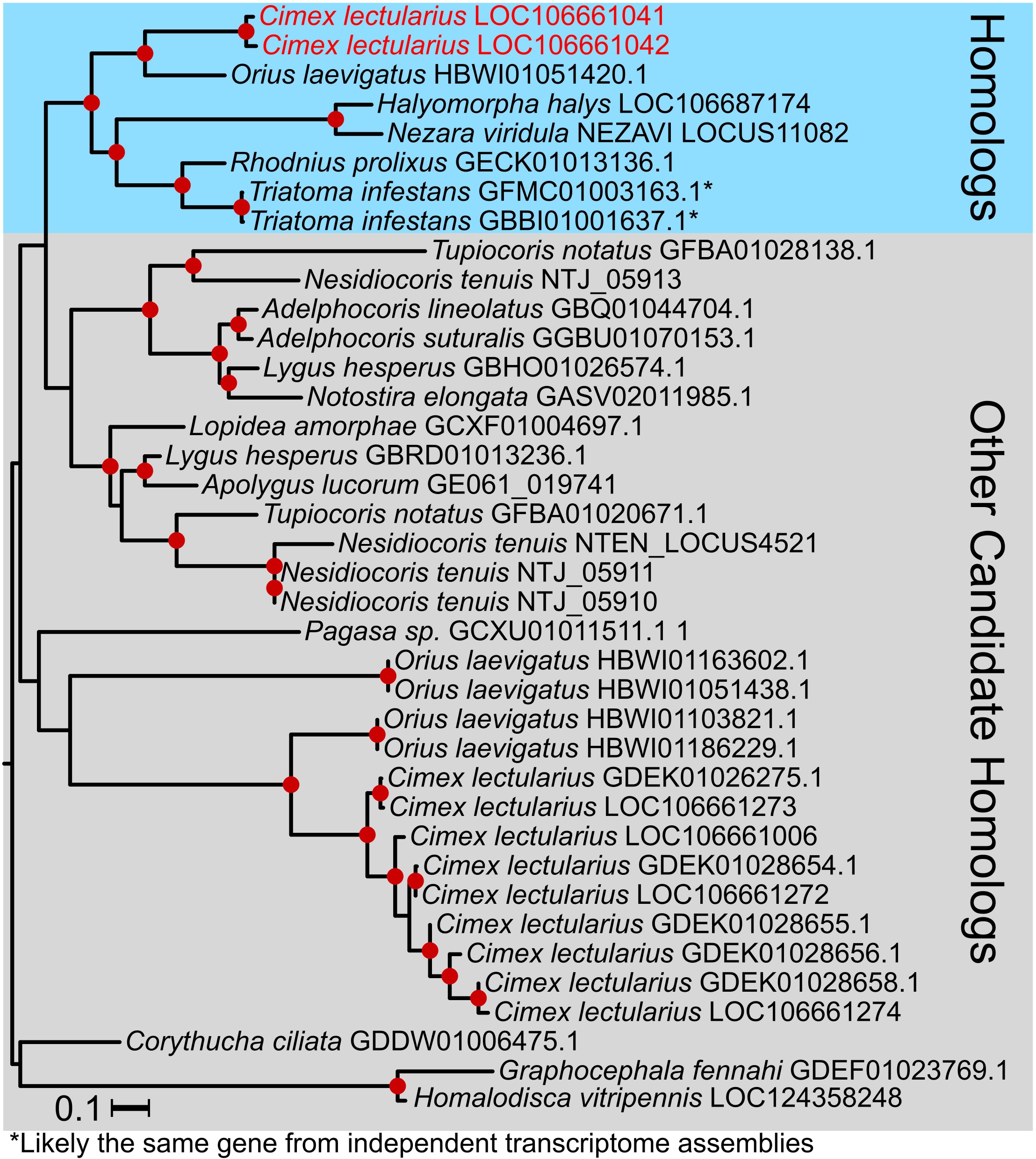
The two most highly expressed genes in response to *P. fluorescens* are paralogous. Red dots along the tree indicate ultrafast bootstrap support ≥ 75. The blue box indicates a clade including the two *C. lectularius* genes of interest (LOC106661041 and LOC106661042-in red text), along with their putative orthologs. The gray box indicates all other genes that were used as candidate homologs for phylogenetic analysis. The asterisks next to the *Triatoma infestans* sequences indicate that they are likely the same gene from different transcriptome assemblies, making *C. lectularius* the only known lineage with a duplication of this gene.

### Infection with traumatic insemination bacteria

When we challenged bed bugs using bacteria cultured from a swab of their enclosure, we found 30 genes upregulated and two genes downregulated (**Fig. 1B**). Similar to the known entomopathogen, this treatment also upregulated immune effector genes, and we found significant enrichment of GO terms describing immune function in the set of differentially expressed genes, indicating detection of these bacteria by the humoral immune system (**Table 2**). This treatment had the least number of genes that were differentially expressed, but of the 30 differentially expressed genes, five of them encode for antimicrobial effectors: Cl-defensin A, Cl-definsin B, Cl-lysozyme, Cl-prolixicin A, and Cl-prolixicin B. Because both Cl-prolixicin genes were upregulated in response to this treatment and no studies have assessed the evolutionary relationship of these two genes, we built a phylogeny with them including *Drosophila melanogaster* diptericin paralogs and a selection of candidate homologs identified from BLAST. We found that the two bed bug prolixicins are likely the result of a gene duplication specific to the bed bug lineage, and that these genes form a clade with prolixicin, an experimentally validated AMP from the kissing bug *Rhodnius prolixus* (**Fig. 3**).

**Fig. 3.**
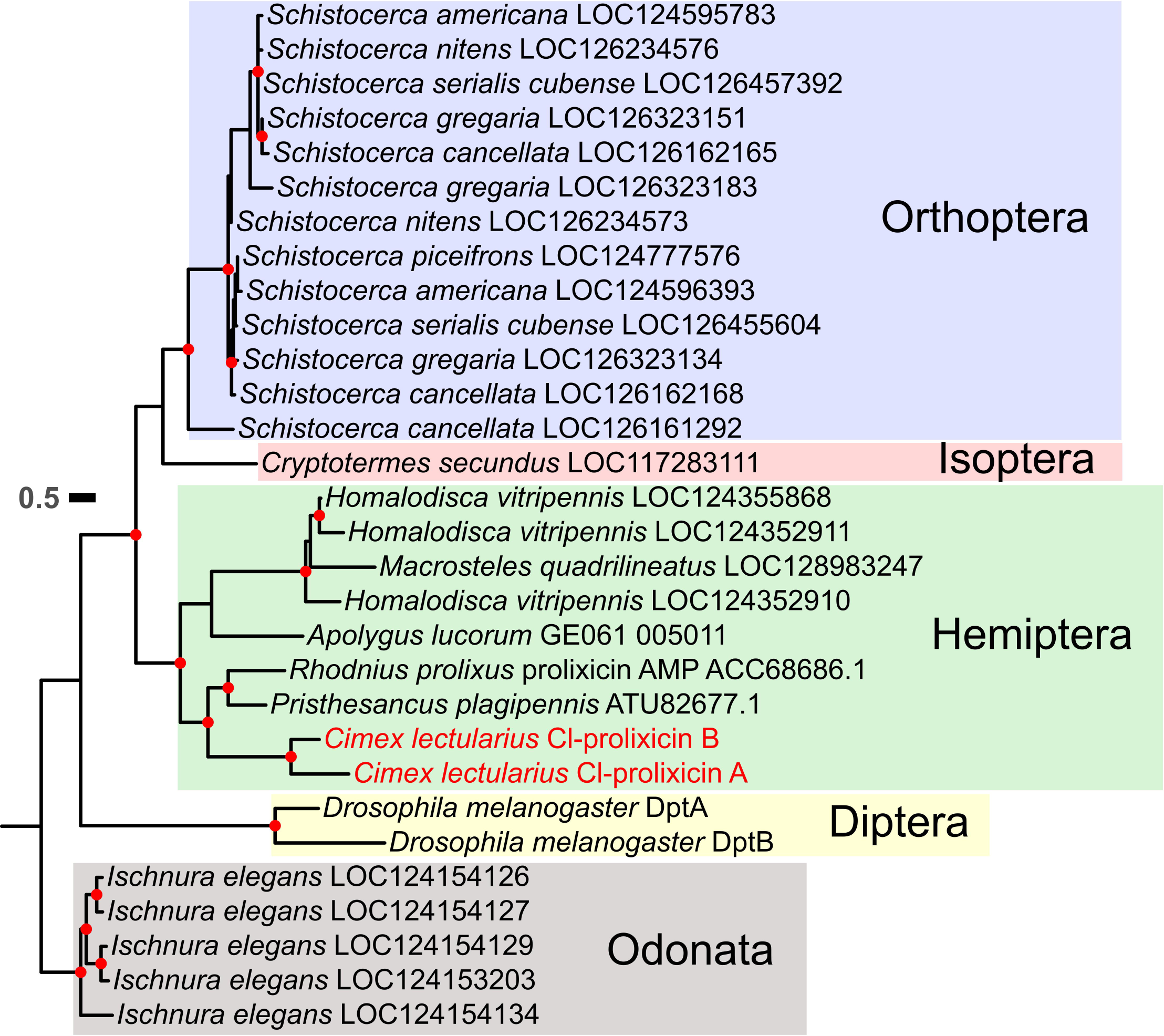
*Cimex lectularius* diptericin-like genes are paralogous and phylogenetically related to the *Rhodnius prolixus* prolixicin AMP. Ultrafast bootstrap values ≥ 75 are represented by red dots along the tree. The clade of genes from the order Odonata was used as the outgroup, as it is the deepest branching lineage in the phylogeny [51].

**Table 2.** Top 10 significantly enriched biological process GO terms found in the genes differentially expressed in response to simulating traumatic insemination.

| GO ID | Term | P value |
| --- | --- | --- |
| GO:0050830 | defense response to Gram-positive bacterium | 0.00021 |
| GO:0050829 | defense response to Gram-negative bacterium | 0.00072 |
| GO:0019731 | antibacterial humoral response | 0.00110 |
| GO:0032365 | intracellular lipid transport | 0.00134 |
| GO:0042742 | defense response to bacterium | 0.00548 |
| GO:0044248 | cellular catabolic process | 0.00578 |
| GO:0044282 | small molecule catabolic process | 0.00861 |
| GO:0006577 | amino-acid betaine metabolic process | 0.00895 |
| GO:0009437 | carnitine metabolic process | 0.00895 |
| GO:0014060 | regulation of epinephrine secretion | 0.00895 |

### Infection with Borrelia duttoni

We found 21 genes upregulated in response to *B. duttoni*, and 55 genes downregulated (**Fig. 1C**). *B. duttoni* is a human pathogen and is the causative agent of tick-borne relapsing fever. Unlike the other treatments, there was not a clear pattern of differential immune gene expression in response to *B. duttoni* (**Table 3**). Interestingly, Cl-prolixicin B was downregulated (log2 fold change = −2.30) in response to a *B. duttoni* contaminated bloodmeal (**Fig. 1C**). Most of the enriched GO terms in this treatment were related to protein localization to the membrane, which could be reflective of a cellular immune response to *B. duttoni* instead of a humoral (**Table 3**).

**Table 3.** Top 10 significantly enriched biological process GO terms found in the genes differentially expressed following a blood containing the human pathogen *B. duttoni*.

| GO ID | Term | P value |
| --- | --- | --- |
| GO:0002181 | Cytoplasmic translation | 8.1e-07 |
| GO:0006614 | SRP-dependent cotranslational protein targeting to membrane | 4.2e-06 |
| GO:0006613 | Cotranslational protein targeting to membrane | 6.4e-06 |
| GO:0045047 | Protein targeting to endoplasmic reticulum (ER) | 1.3e-05 |
| GO:0072599 | Establishment of protein localization to ER | 2.1e-05 |
| GO:0000184 | Nuclear-transcribed mRNA catabolic process | 2.1e-05 |
| GO:0070972 | protein localization to ER | 2.1e-05 |
| GO:0006612 | Protein targeting to membrane | 0.00014 |
| GO:0090150 | Establishment of protein localization to membrane | 0.00028 |
| GO:0006413 | Translational initiation | 0.00033 |

### Shared responses between treatments

Overall, there are very few overlapping genes between each dataset, with nine overlapping genes differentially expressed in response to the TI bacteria and *P. fluorescens* treatments (Fisher’s exact test, *P* < 0.0001, OR = 62.1, 95% CI = 28.2-Inf), three overlapping in response to the *P. fluorescens* and *B. duttoni* treatments (Fisher’s exact test, *P* =0.02, OR = 5.9, 95% CI = 5.9-Inf), one gene overlapping between the TI bacterial treatment and the *B. duttoni* treatment (Fisher’s exact test, *P* = 0.2, OR = 4.8, 95% CI = 0.2-Inf), and zero shared between all treatments (**Fig. 4**). Intriguingly, the only gene shared between the TI bacterial treatment and the *B. duttoni* treatment was Cl-prolixicin B which was upregulated in response to TI bacteria, but downregulated in response to a *B. duttoni* contaminated blood meal (**Fig. 1B and C**). The three genes that are differentially expressed in both the *B. duttoni* and *P. fluorescens* datasets encode for proteins that are putatively constituents of the insect cuticle (**Fig. 4**). Out of the nine genes that are differentially expressed in both the TI bacteria and the *P. fluorescens* treatments, four of them are antimicrobial immune effectors, while the other five consist of 3 cuticular proteins a laccase-like gene, and a mucin-like gene (**Fig. 4**). Overall, our results suggest that *P. fluorescens* and TI bacteria are detected by the bed bug humoral immune system, while *B. duttoni* ingested through a blood meal may only induce the cellular immune response.

**Fig. 4.**
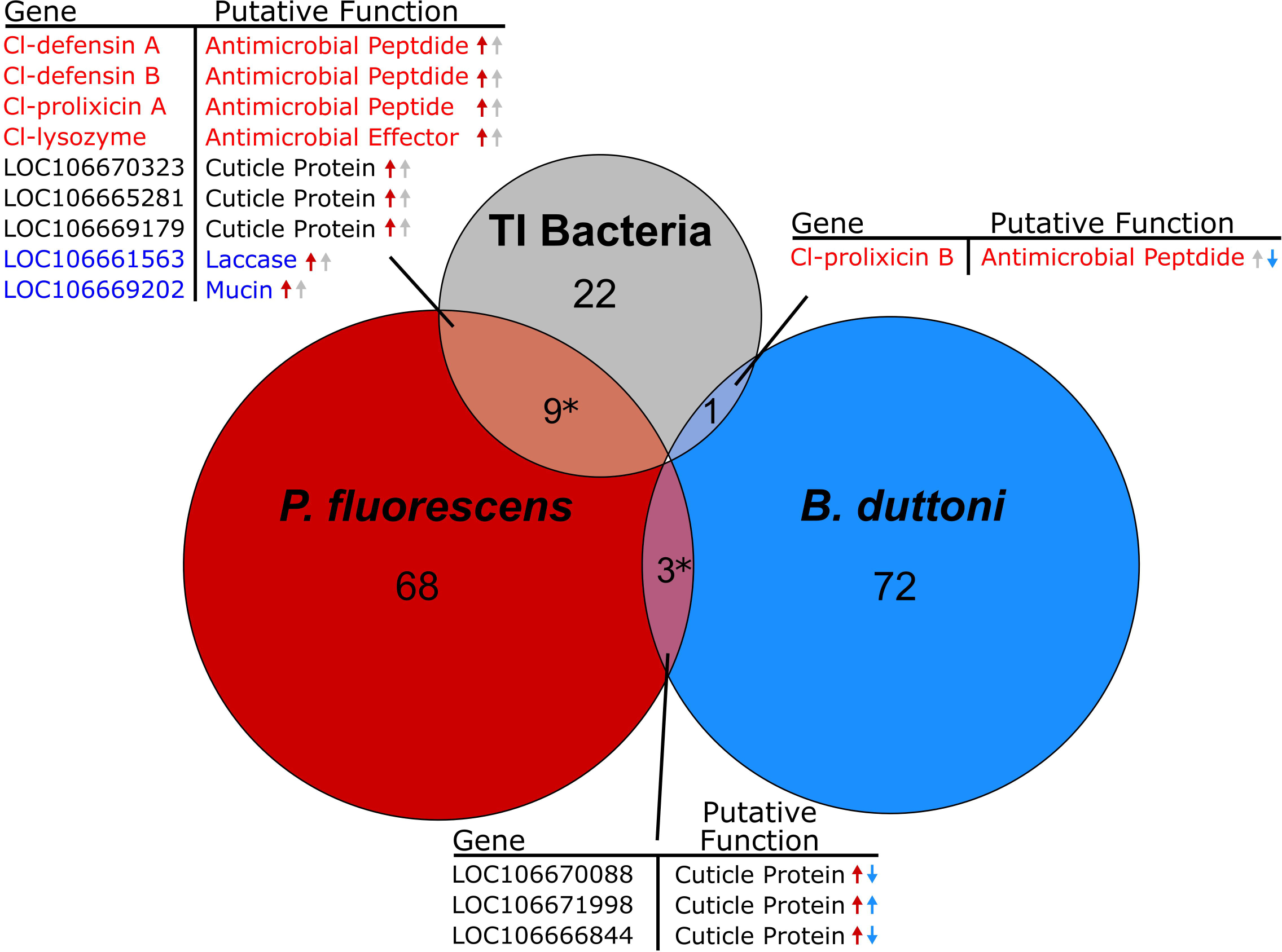
Venn diagram highlighting the distinct transcriptomic response to each treatment. The differentially expressed genes shared between each treatment and their putative functions are detailed in the tables corresponding to the Venn diagram overlaps. The arrows in the tables detail if each gene was upregulated or downregulated in response to its treatment, with the treatment colors corresponding to the circles in the Venn diagram. Genes in red text encode for known immune effectors, and genes in blue text encode for proteins that play immunological roles in other insects. Asterisks indicate significantly more overlap between differentially expressed genes than one would expect at random.

## Discussion

Characterizing the bed bug immune response could help us understand why they are poor vectors while also informing the optimization of biological control approaches that use bacteria [53]. Thus, we exposed bed bugs to various bacterial challenges to further understand their molecular immune response. Interestingly, immune effector genes were differentially upregulated in the *P. fluorescens* and TI bacterial treatments, but there were no upregulated immune genes in response to a blood meal contaminated with *B. dutton*i. This suggests that there is *bona fide* molecular immune detection of *P. flourescens* and TI bacteria, but *B. duttoni* may go undetected by bed bug PRRs in the gut.

The annotated immune effectors in the bed bug genome that were differentially expressed in our study were two of the three *C. lectularius* defensins, one of four lysozyme-like genes, and both *C. lectularius* diptericin-like (prolixicin) genes. All of these were upregulated in response to both *P. fluorescens* and the TI bacterial challenge except for one diptericin-like gene, Cl-prolixicin B, which was only upregulated in response to the TI bacteria (**Fig. 1B**). This could mean that the bed bug diptericin-like AMPs have specific responses to different bacterial taxa, as the *P. fluorescens* treatment consisted of a single species of bacteria, while the TI bacterial treatment probably included a variety of species. Interestingly, *Drosophila melanogaster* diptericins have differential action against bacteria, with one having specific antimicrobial activity against *Providencia rettgeri*, and the other having antimicrobial activity against *Acetobacter* species [54]. Furthermore, the retention of each diptericin gene copy was correlated to *Drosophila* ecology, with some species losing a paralog depending on the microbial community associated with their food source [54]. Because both Cl-prolixicin genes were upregulated in response to the TI bacteria, but only Cl-prolixicin A was upregulated in response to *P. fluorescens*, Cl-prolixicin B may be an important effector in response to wounding by traumatic insemination. This is supported by a study of the bed bug spermalege (the receptacle organ for traumatic insemination) transcriptome, where a diptericin-like bed bug transcript was highly represented [19].

In the *B. duttoni* treatment, Cl-prolixicin B was downregulated, and no other immune effectors were differentially expressed, suggesting that *B. duttoni* is not detected by the bed bug humoral immune system and is either 1.) cleared by other means or 2.) persists temporally. A previous study found that another *Borrelia* species, *Borrelia recurrentis*, interacts with extracellular DNA traps, a cellular response to pathogen infection carried out by hemocytes in culture [32]. The use of extracellular DNA traps as a pathogen defense is a relatively new finding in insects [17,55–58]. They are made of extracellular chromatin and proteins to trap pathogens so that hemocytes can engulf them [32,57,58]. Interestingly, two histone-like genes (LOC106672213 and LOC106663736) were downregulated in response to *B. duttoni* infection, and a peroxidase-like gene (LOC106661969) was upregulated. Both histones and peroxidases are known constituents of extracellular DNA traps in other systems [57,59–61]. Using reciprocal BLAST searches, we found that the peroxidase-like gene (LOC106661969) upregulated in response to *B. duttoni* infection is homologous to peroxinectin, which is a constituent of extracellular DNA traps in the crustacean *Carcinus maenas* [61]. These results suggest that *Borrelia* species may evade pathogen recognition by the humoral immune system of bed bugs, but could be cleared by cellular immune responses [29,32,33].

Interestingly, many genes encoding for proteins putatively involved in cuticle formation were differentially expressed in response to our three experimental treatments, especially the *P. fluorescens* treatment (**Table 1**). Not only is the insect cuticle involved in immunity as the first line of defense to pathogens as a physical barrier, but there is some overlap in cuticular formation pathways and insect humoral immunity [22]. The best example of this is the phenoloxidase pathway, which is involved in both melanization/sclerotization of the cuticle and participates in the insect humoral immune response as it encapsulates invading pathogens by melanization [21,22]. Two types of phenoloxidase enzymes are proposed to be responsible for sclerotization: tyrosinases and laccases. A multicopper oxidase domain containing gene (LOC106661563) annotated as laccase-5 was differentially upregulated in the *P. fluorescens* and the TI bacteria treatments (**Fig. 4**). Laccase genes have been shown to be involved in cuticle tanning and immunity in various arthropods, and knockdown of laccase genes has led to increases in pathogen susceptibility in beetles and mosquitoes [62–67]. LOC106661563 has never been considered in bed bug immune studies but it could be an interesting candidate for functional studies. We did not find the previously annotated bed bug phenoloxidase effector genes (LOC106673939 and LOC106673564) differentially expressed in our treatments, but they are constitutively expressed in all samples (**Additional File 1: Fig. S1**). Differences in expression of prophenoloxidases at the transcriptomic level may not be easily detected because these proteins exist as inactive precursors, or zymogens. The immunological role of cuticle proteins in in bed bugs could be a worthy avenue of research as their cuticle likely faced extreme selection pressures for elasticity from blood feeding and traumatic insemination, and potentially wound healing and pathogen clearance during traumatic insemination. In further support, cuticular genes were differentially expressed in every treatment administered in this study, and some were differentially expressed in multiple treatments **(Fig. 4**).

Another gene that was differentially expressed in both the *P. fluorescens* treatment, and the TI bacterial treatment is a mucin-2-like gene (**Fig.** 4). Mucin proteins help form the mucosal linings of some insect tissues, and they can also have an immunological component as they serve as a line of defense against pathogens [70,71]. Furthermore, one study found that a mucin protein may be involved in bacterial entrapment and immunity in *Drosophila* [72]. The mucin-like gene that was differentially expressed in our study encodes 10 protein isoforms between 1698-2021 amino acids in length. It contains multiple conserved protein domains including a zona pellucida domain (cl27758), an atrophin-1 family domain (cl26464), and a topoisomerase II-associated protein PAT1 domain (cl25764). Although the actual function of this mucin-like gene is unknown, it could be a candidate for future functional assessment of immune insult in bed bugs.

The two most differentially expressed genes of any treatment in this study were two paralogs (**Fig. 2**) that were highly upregulated in response to *P. fluorescens* treatment. One of the genes (LOC106661042) is described as “acanthoscurrin-like” in the NCBI database, while the other (LOC106661041) is described as “glycine rich protein DOT-1-like.” Acanthoscurrin is an antimicrobial peptide that was discovered in a species of tarantula [52]. However, upon further investigation, LOC106661042 has very little similarity to acanthoscurrin. Instead, we suspect that LOC106661041 and LOC106661042 are more likely to be cuticular proteins because of their high glycine content (∼40%) [14]. Interestingly, LOC106661041 and LOC106661042 belong to a highly supported clade of genes from only heteropteran insects, while the nodes leading up to it are all low support (**Fig. 2**). This suggests that the rest of the genes included in the phylogeny are distant homologs, and that LOC106661041 and LOC106661042 belong to a gene family that is novel to the Heteroptera (**Fig. 2**). None of the other genes in this clade are characterized, giving us no further insight to their possible function. Although LOC106661041 and LOC106661042 may not be AMPs, their role in bed bug immunity should still be assessed.

## Conclusions

Our study opens some interesting questions about the response of bed bugs to immune challenges. First, there is substantial overlap between the immune transcriptomic response between a known bed bug pathogen and the bacteria that are likely encountered during traumatic insemination (**Fig. 4**). Both treatments induced the upregulation of immune effectors in bed bugs, providing clear evidence of molecular recognition of microbes. Upregulation of immune effectors by the treatment of TI associated microbes supports previous findings suggesting that the bacteria encountered during traumatic insemination cause a *bona fide* immune insult [11,18,69]. Second, a bloodmeal contaminated with a human pathogen did not upregulate any of known AMP effectors in bed bugs, and alternatively, downregulated the only immune effector that was differentially expressed in this treatment (**Fig. 1C**). This suggests that *B. duttoni* is not detected by the humoral immune system in bed bugs and is either cleared by the cellular immune responses or persists undetected. Alternatively, *B. duttoni* could activate effectors that are hard to detect using transcriptomics. Still, our study warrants more investigation of the bed bug immune response to *Borrelia* spp. and other vector-borne pathogens given the differences we observed. Finally, we identified multiple candidate genes for future studies of the bed bug immune system, including putative cuticle-related proteins, a laccase-like gene, and a mucin-like gene.

## Declarations

### Acknowledgements

The graphical abstract was created with BioRender.com.

### Availability of data and materials

All raw sequencing datasets generated in this study are available in NCBI’s Sequence Read Archive (SRA) under the BioProject Accession: PRJNA1079782.

### Authors’ contributions

H.K.W performed and oversaw all data analysis, wrote the manuscript. A.B. performed the TI bacteria laboratory experiments. S.E.M. performed the differential expression analysis and gene ontology enrichment analysis. J.A.H. retrieved sequence data and performed the phylogenetic analysis of the acanthoscurrin-like genes and the glycine rich dot-1-like genes. J.G..K was involved in the conceptualization of the study, supervised HKW and A.B., and acquired funding for sequencing. F.G.H. supervised H.K.W., S.E.M., and J.A.H. throughout all computational analyses. JEP was involved in the conceptualization of the study, acquired funding for the laboratory experiments, performed the *P. fluorescens* and *B. duttoni* experiments. All authors reviewed the manuscript.

### Ethics approval and consent to participate

Not applicable

### Consent for publication

Not applicable

## References

1. Goddard J, De Shazo R. Psychological effects of bed bug attacks (Cimex lectularius L.). Am J Med. 2012;125:101–3. doi:10.1016/j.amjmed.2011.08.010

2. Goddard J, deShazo R. Bed bugs (*Cimex lectularius*) and clinical consequences of their bites. JAMA 2009;301:1358–66. doi:10.1001/jama.2009.405

3. Ho D, Lai O, Glick S, Jagdeo J. Lack of evidence that bedbugs transmit pathogens to humans. J Am Acad Dermatol. 2016;74:1261. doi:10.1016/j.jaad.2015.12.007

4. Lai O, Ho D, Glick S, Jagdeo J. Bed bugs and possible transmission of human pathogens: a systematic review. Arch Dermatol Res. 2016;308:531–8. 10.1007/s00403-016-1661-8

5. Pietri JE. Case not closed: Arguments for new studies of the interactions between bed bugs and human pathogens. Am J Trop Med Hyg. 2020;103:619–24.

6. Fisher ML, Levine JF, Guy JS, Mochizuki H, Breen M, Schal C, et al. Lack of influence by endosymbiont *Wolbachia* on virus titer in the common bed bug, Cimex lectularius. Parasit Vectors. 2019;12:436.

7. Delaunay P, Blanc V, Del Giudice P, Levy-Bencheton A, Chosidow O, Marty P, et al. Bedbugs and infectious diseases. Clin Infect Dis. 2011;52:200–10.

8. Usinger RL. Monograph of Cimicidae (Hemiptera, Heteroptera). Entomological Society of America; 1966.

9. Morrow EH, Arnqvist G. Costly traumatic insemination and a female counter-adaptation in bed bugs. Proc R Soc B Biol Sci. 2003;270:2377–81.

10. Stutt AD, Siva-Jothy MT. Traumatic insemination and sexual conflict in the bed bug *Cimex lectularius*. Proc Natl Acad Sci U S A. 2001;98:5683–7.

11. Reinhardt K, Naylor R, Siva-Jothy MT. Reducing a cost of traumatic Insemination: female bedbugs evolve a unique organ. Proc Biol Sci. 2003;270:2371–5. doi: 10.1098/rspb.2003.2515

12. Roth S, Siva-Jothy MT, Balvín O, Morrow EH, Willassen E, Reinhardt K. The evolution of female-biased genital diversity in bedbugs (Cimicidae). Evolution. 2023;qpad211. doi:10.1093/evolut/qpad211

13. Michels J, Gorb SN, Reinhardt K. Reduction of female copulatory damage by resilin represents evidence for tolerance in sexual conflict. J R Soc Interface. 2015;12:20141107.

14. Benoit JB, Adelman ZN, Reinhardt K, Dolan A, Poelchau M, Jennings EC, et al. Unique features of a global human ectoparasite identified through sequencing of the bed bug genome. Nat Commun. 2016;7:1–10.

15. Cho Y, Cho S. Hemocyte-hemocyte adhesion by granulocytes is associated with cellular immunity in the cricket, Gryllus bimaculatus. Sci Rep. 2019;9:18066. doi:10.1038/s41598-019-54484-5

16. Eleftherianos I, Heryanto C, Bassal T, Zhang W, Tettamanti G, Mohamed A. Haemocyte-mediated immunity in insects: Cells, processes and associated components in the fight against pathogens and parasites. Immunology. 2021;164:401–32.

17. Potts R, King JG, Pietri JE. Ex vivo characterization of the circulating hemocytes of bed bugs and their responses to bacterial exposure. J Invertebr Pathol. 2020;174:107422.

18. Siva-Jothy MT, Zhong W, Naylor R, Heaton L, Hentley W, Harney E. Female bed bugs (Cimex lectularius L) anticipate the immunological consequences of traumatic insemination via feeding cues. Proc Natl Acad Sci U S A. 2019;116:14682–7.

19. Moriyama M, Koga R, Hosokawa T, Nikoh N, Futahashi R, Fukatsu T. Comparative transcriptomics of the bacteriome and the spermalege of the bedbug Cimex lectularius (Hemiptera: Cimicidae). Appl Entomol Zool. 2012;47:233–43. doi:10.1007/s13355-012-0112-z

20. Pietri JE, Liang D. Virulence of entomopathogenic bacteria in the bed bug, Cimex lectularius. J Invertebr Pathol. 2018;151:1–6.

21. Hillyer JF. Insect immunology and hematopoiesis. Dev Comp Immunol. 2016;58:102–18.

22. Siva-Jothy MT, Moret Y, Rolff J. Insect immunity: an evolutionary ecology perspective. In: Simpson SJBT-A in IP, editor. Academic Press; 2005. p. 1–48.

23. Zumaya-Estrada FA, Martínez-Barnetche J, Lavore A, Rivera-Pomar R, Rodríguez MH. Comparative genomics analysis of triatomines reveals common first line and inducible immunity-related genes and the absence of Imd canonical components among hemimetabolous arthropods. Parasit Vectors. 2018;11:48. doi:10.1186/s13071-017-2561-2

24. Meraj S, Dhari AS, Mohr E, Lowenberger C, Gries G. Characterization of new defensin antimicrobial peptides and their expression in bed bugs in response to bacterial ingestion and injection. Int J Mol Sci. 2022;23.

25. Meraj S, Mohr E, Ketabchi N, Bogdanovic A, Lowenberger C, Gries G. Time- and tissue-specific antimicrobial activity of the common bed bug in response to blood feeding and immune activation by bacterial injection. J Insect Physiol. 2021;135:104322. doi:10.1016/j.jinsphys.2021.104322

26. Bellinvia S, Johnston PR, Mbedi S, Otti O. Mating changes the genital microbiome in both sexes of the common bedbug *Cimex lectularius* across populations. Proc Biol. Sci 2020;287:20200302. doi: 10.1098/rspb.2020.0302

27. Kaushal A, Gupta K, van Hoek ML. Characterization of *Cimex lectularius* (bedbug) defensin peptide and its antimicrobial activity against human skin microflora. Biochem Biophys Res Commun. 2016;470:955–60. doi10.1016/j.bbrc.2016.01.100

28. Leulmi H, Bitam I, Berenger JM, Lepidi H, Rolain JM, Almeras L, et al. Competence of *Cimex lectularius* bed bugs for the transmission of *Bartonella quintana*, the agent of trench fever. PLoS Negl Trop Dis. 2015;9:4–15.

29. El Hamzaoui B, Laroche M, Bechah Y, Bérenger JM, Parola P. Testing the competence of *Cimex lectularius* bed bugs for the transmission of *Borrelia recurrentis*, the agent of relapsing fever. Am J Trop Med Hyg. 2019;100:1407–12.

30. Salazar R, Castillo-Neyra R, Tustin AW, Borrini-Mayorí K, Náquira C, Levy MZ. Bed bugs (*Cimex lectularius*) as vectors of *Trypanosoma cruzi*. Am J Trop Med Hyg. 2015;92:331–5.

31. Blakely BN, Hanson SF, Romero A. Survival and Transstadial Persistence of Trypanosoma cruzi in the bed bug (Hemiptera: Cimicidae). J Med Entomol. 2018;55:742–6.

32. Potts R, Scholl JL, Baugh LA, Pietri JE. A comparative study of body lice and bed bugs reveals factors potentially involved in differential vector competence for the relapsing fever spirochete *Borrelia recurrentis*. Infect Immun. 2022;90:e0068321.

33. Blanc G, Bruneau J, Chabaud A. The behaviour of certain spirochaetes in *Cimex lectularius*. Arch Inst Pasteur Maroc. 1953;4:411–28.

34. Andrews S. FASTQC. A quality control tool for high throughput sequence data. 2010.

35. Bolger AM, Lohse M, Usadel B. Trimmomatic: A flexible trimmer for Illumina sequence data. Bioinformatics. 2014;30:2114–20.

36. Patro R, Duggal G, Love MI, Irizarry RA, Kingsford C. Salmon provides fast and bias-aware quantification of transcript expression. Nat Methods. 2017;14:417–9. doi:10.1038/nmeth.4197

37. Soneson C, Love MI, Robinson MD. Differential analyses for RNA-seq: transcript-level estimates improve gene-level inferences. F1000Research. 2015;4:1521.

38. Love MI, Huber W, Anders S. Moderated estimation of fold change and dispersion for RNA-seq data with DESeq2. Genome Biol. 2014;15:1–21.

39. Cantalapiedra CP, Hernández-Plaza A, Letunic I, Bork P, Huerta-Cepas J. eggNOG-mapper v2: Functional Annotation, Orthology Assignments, and Domain Prediction at the Metagenomic Scale. Mol Biol Evol. 2021;38:5825–9. doi:10.1093/molbev/msab293

40. Buchfink B, Xie C, Huson DH. Fast and sensitive protein alignment using DIAMOND. Nat Methods. 2015;12:59–60. doi:10.1038/nmeth.3176

41. Buchfink B, Reuter K, Drost H-G. Sensitive protein alignments at tree-of-life scale using DIAMOND. Nat Methods. 2021;18:366–8. doi:10.1038/s41592-021-01101-x

42. Alexa A, Rahnenfuhrer J. topGO: Enrichment Analysis for Gene Ontology. 2020.

43. R Core Team. R: A Language and Environment for Statistical Computing [Internet]. Vienna, Austria; 2020. https://www.r-project.org/

44. Suyama M, Torrents D, Bork P. PAL2NAL: robust conversion of protein sequence alignments into the corresponding codon alignments. Nucleic Acids Res. 2006;34:W609–12. doi:10.1093/nar/gkl315

45. Minh BQ, Schmidt HA, Chernomor O, Schrempf D, Woodhams MD, von Haeseler A, et al. IQ-TREE 2: new models and efficient methods for phylogenetic inference in the genomic era. Mol Biol Evol. 2020;37:1530–4. doi:10.1093/molbev/msaa015

46. Kalyaanamoorthy S, Minh BQ, Wong TKF, von Haeseler A, Jermiin LS. ModelFinder: fast model selection for accurate phylogenetic estimates. Nat Methods. 2017;14:587–9. doi:10.1038/nmeth.4285

47. Anisimova M, Gil M, Dufayard JF, Dessimoz C, Gascuel O. Survey of branch support methods demonstrates accuracy, power, and robustness of fast likelihood-based approximation schemes. Syst Biol. 2011;60:685–99.

48. Minh BQ, Nguyen MAT, Von Haeseler A. Ultrafast approximation for phylogenetic bootstrap. Mol Biol Evol. 2013;30:1188–95.

49. Nguyen L-T, Schmidt HA, von Haeseler A, Minh BQ. IQ-TREE: A fast and effective stochastic algorithm for estimating maximum-likelihood phylogenies. Mol Biol Evol. 2015;32:268–74. doi: 10.1093/molbev/msu300

50. Hoang DT, Chernomor O, von Haeseler A, Minh BQ, Vinh LS. UFBoot2: Improving the ultrafast bootstrap approximation. Mol Biol Evol. 2018;35:518–22. doi: 10.1093/molbev/msx281

51. Misof B, Liu S, Meusemann K, Peters RS, Donath A, Mayer C, et al. Phylogenomics resolves the timing and pattern of insect evolution. Science. 2014;346:763–7. doi:10.1126/science.1257570

52. Lorenzini DM, da Silva PIJ, Fogaça AC, Bulet P, Daffre S. Acanthoscurrin: a novel glycine-rich antimicrobial peptide constitutively expressed in the hemocytes of the spider *Acanthoscurria gomesiana*. Dev Comp Immunol. 2003;27:781–91.

53. Pietri JE, Potts R. Effects of NF-kB signaling inhibitors on bed bug resistance to orally provisioned entomopathogenic bacteria. Insects. 2021;12.

54. Hanson MA, Grollmus L, Lemaitre B. Ecology-relevant bacteria drive the evolution of host antimicrobial peptides in *Drosophila*. Science. 2023;381:eadg5725. doi: 10.1126/science.adg5725

55. Coste Grahl M V, Perin APA, Lopes FC, Porto BN, Uberti AF, Canavoso LE, et al. The role of extracellular nucleic acids in the immune system modulation of Rhodnius prolixus (Hemiptera: Reduviidae). Pestic Biochem Physiol. 2020;167:104591. doi: 10.1016/j.pestbp.2020.104591

56. Chen RY, Keddie BA. *Galleria mellonella* (Lepidoptera: Pyralidae) hemocytes release extracellular traps that confer protection against bacterial infection in the hemocoel. J Insect Sci. 2021;21:17. doi:10.1093/jisesa/ieab092

57. Nascimento MTC, Silva KP, Garcia MCF, Medeiros MN, Machado EA, Nascimento SB, et al. DNA extracellular traps are part of the immune repertoire of *Periplaneta americana*. Dev Comp Immunol. 2018;84:62–70. doi: 10.1016/j.dci.2018.01.012

58. Altincicek B, StoLJtzel S, Wygrecka M, Preissner KT, Vilcinskas A. Host-derived extracellular nucleic acids enhance innate immune responses, induce coagulation, and prolong survival upon infection in insects. J Immunol. 2008;181:2705–12.

59. Brinkmann V, Reichard U, Goosmann C, Fauler B, Uhlemann Y, Weiss DS, et al. Neutrophil extracellular traps kill bacteria. Science. 2004;303:1532–5.

60. Mamtimin M, Pinarci A, Han C, Braun A, Anders HJ, Gudermann T, et al. Extracellular DNA traps: origin, function and implications for anti-cancer therapies. Front Oncol. 2022;12:1– 34.

61. Robb CT, Dyrynda EA, Gray RD, Rossi AG, Smith VJ. Invertebrate extracellular phagocyte traps show that chromatin is an ancient defence weapon. Nat Commun. 2014;5.

62. Arakane Y, Muthukrishnan S, Beeman RW, Kanost MR, Kramer KJ. Laccase 2 is the phenoloxidase gene required for beetle cuticle tanning. Proc Natl Acad Sci. 2005;102:11337–42. doi:10.1073/pnas.0504982102

63. Du M-H, Yan Z-W, Hao Y-J, Yan Z-T, Si F-L, Chen B, et al. Suppression of laccase 2 severely impairs cuticle tanning and pathogen resistance during the pupal metamorphosis of *Anopheles sinensis* (Diptera: Culicidae). Parasit Vectors. 2017;10:171. doi:10.1186/s13071-017-2118-4

64. Chen Y-H, Song F, Miao Y-T, He H-H, Lian Y-Y, Li X, et al. A novel laccase gene from *Litopenaeus vannamei* is involved in the immune responses to pathogen infection and oxidative stress. Dev Comp Immunol. 2020;105:103582. doi:10.1016/j.dci.2019.103582

65. Shi L, Chan S, Li C, Zhang S. Identification and characterization of a laccase from *Litopenaeus vannamei* involved in anti-bacterial host defense. Fish Shellfish Immunol. 2017;66:1–10. doi: 10.1016/j.fsi.2017.04.026

66. Wang Z, Hu R, Ye X, Huang J, Chen X, Shi M. Laccase 1 gene from *Plutella xylostella* (PxLac1) and its functions in humoral immune response. J Insect Physiol. 2018;107:197–203. doi: 10.1016/j.jinsphys.2018.04.001

67. Hayakawa Y, Sawada M, Seki M, Sirasoonthorn P, Shiga S, Kamiya K, et al. Involvement of laccase2 and yellow-e genes in antifungal host defense of the model beetle, *Tribolium castaneum*. J Invertebr Pathol. 2018;151:41–9. doi: 10.1016/j.jip.2017.10.010

68. An C, Budd A, Kanost MR, Michel K. Characterization of a regulatory unit that controls melanization and affects longevity of mosquitoes. Cell Mol Life Sci. 2011;68:1929–39.

69. Reinhardt K, Naylor RA, Siva-Jothy MT. Potential sexual transmission of environmental microbes in a traumatically inseminating insect. Ecol Entomol. 2005;30:607–11.

70. Syed ZA, Härd T, Uv A, van Dijk-Härd IF. A potential role for *Drosophila* mucins in development and physiology. PLoS One. 2008;3:e3041.

71. Zeng T, Jaffar S, Xu Y, Qi Y. The intestinal immune defense system in Insects. Int J Mol Sci. 2022;23.

72. Korayem AM, Fabbri M, Takahashi K, Scherfer C, Lindgren M, Schmidt O, et al. A *Drosophila* salivary gland mucin is also expressed in immune tissues: evidence for a function in coagulation and the entrapment of bacteria. Insect Biochem Mol Biol. 2004;34:1297–304. doi:10.1016/j.ibmb.2004.09.001

