## Additional File 1 for "Under the radar: differential responses of bed bugs to an entomopathogen, environmental bacteria, and a human pathogen" (from another version of this preprint: version 1, 2024-03-01)

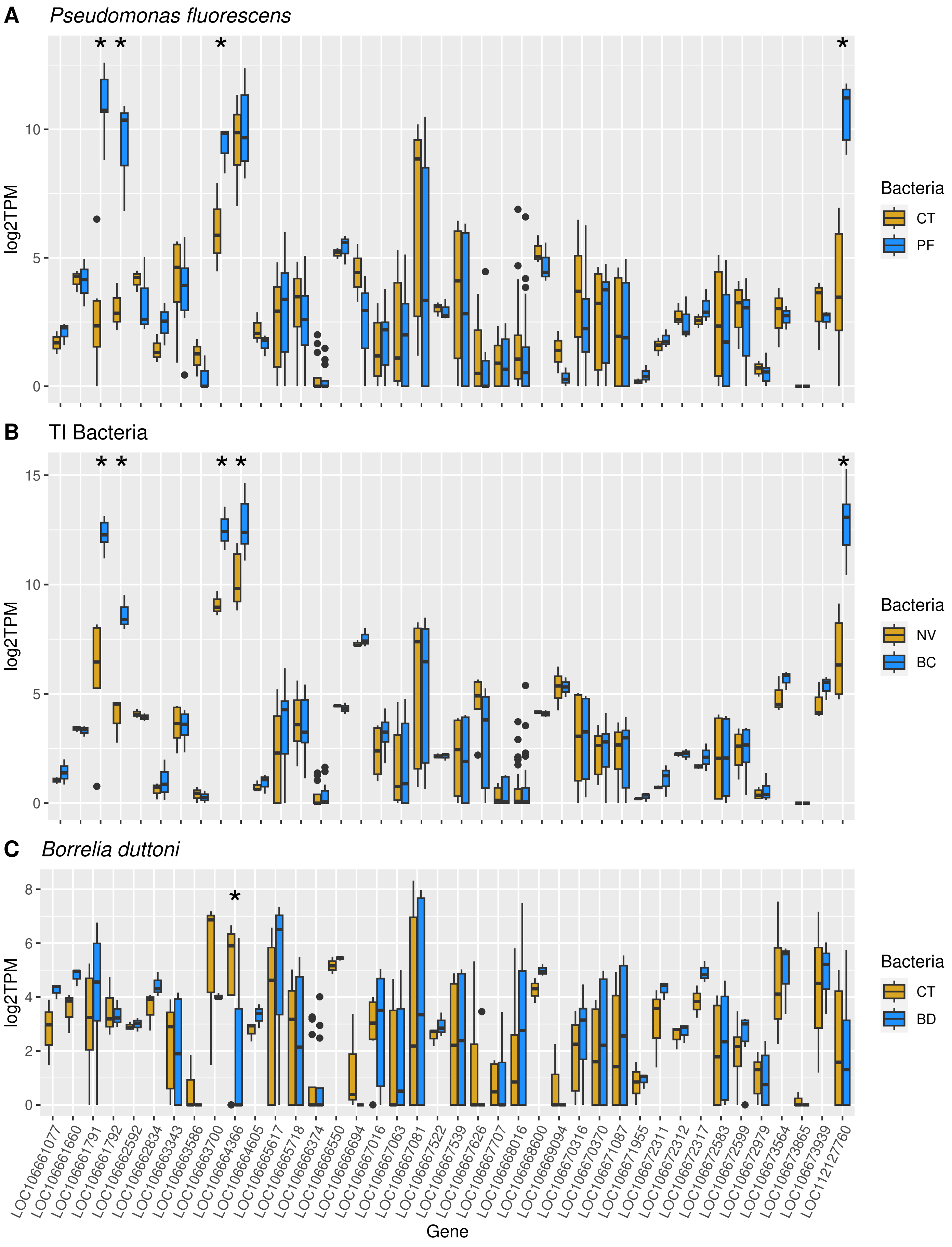
A
Pseudomonas fluorescens
log2TPM
0
5
10
Bacteria
CT
PF
B
TI Bacteria
log2TPM
0
5
10
15
Bacteria
NV
BC
C
Borrelia duttoni
log2TPM
0
2
4
6
8
Bacteria
CT
BD
LOC106661077
LOC106661660
LOC106661791
LOC106661792
LOC106662592
LOC106662834
LOC106663343
LOC106663586
LOC106663700
LOC106664366
LOC106664605
LOC106665617
LOC106665718
LOC106666374
LOC106666550
LOC106666694
LOC106667016
LOC106667063
LOC106667081
LOC106667522
LOC106667539
LOC106667626
LOC106667707
LOC106668016
LOC106668600
LOC106669094
LOC106670316
LOC106670370
LOC106671087
LOC106671955
LOC106672311
LOC106672312
LOC106672317
LOC106672583
LOC106672599
LOC106672979
LOC106673564
LOC106673865
LOC106673939
LOC112127760
Gene

**Figure S1**: **Immune-related gene expression across all experimental groups**. All TPM values obtained from Salmon were added by one and $log_2$ transformed. Asterisks indicate significant differential expression from previous DEseq2 analysis. CT: Control; PF: *Pseudomonas fluorescens*, NV: unwounded; BC: Bacteria; BD: *Borrelia duttoni*
